# Flexible information coding in frontoparietal cortex across the functional stages of cognitive processing

**DOI:** 10.1101/246132

**Authors:** Joyce M. G. Vromen, Stefanie I. Becker, Oliver Baumann, Jason B. Mattingley, Roger W. Remington

## Abstract

Neural activity in frontoparietal cortex shows overlap across cognitive domains and has been proposed to reflect flexible information processing according to current task demands (Dosenbach et al., 2007; Duncan, 2001). However, a strong assertion of flexibility requires investigating activity across stages of cognitive processing. The current study assessed neural activity in Multiple Demand (MD) regions across the stages of processing that form the core of long-standing cognitive models (Welford, 1952). Specifically, many complex tasks share a comparable structure of subsequent operations: target selection, stimulus-response (SR) mapping, and response execution. We independently manipulated the difficulty of target selection and SR mapping in identical stimulus displays and assessed changes in frontoparietal activity with increased demands in either stage. The results confirmed flexibility in MD regions, with enhanced information representation during difficult target selection as well as SR mapping. Additionally, anterior insula (AI) and anterior cingulate cortex (ACC) showed preferential representation of SR stage information, whereas the medial frontal gyrus (MFG) and inferior parietal sulcus (IPS) showed preferential representation of target selection-stage information. Together these results suggest that MD regions dynamically alter the information they represent with changing task demands. This is the first study to demonstrate that MD regions support flexible goal-directed cognition across multiple processing stages. At the same time we show a preference for the representation of information from a specific processing stage in a subset of MD regions.

**Significance Statement:** Goal-directed cognition in complex tasks is critical to key life outcomes including longevity and academic performance. Nevertheless, the mechanisms underlying cognition in complex tasks are not well understood. Distinct neural networks are critical to the navigation of specific cognitive domains (e.g. attention), but frontoparietal activity shows cross-domain and -task overlap and supports flexible representation of goal-critical information. This study links flexible frontoparietal processing to longstanding models of meta-cognition that propose a unifying structure of operations underlying most tasks: target selection, SR mapping, and response execution. Our results demonstrate that flexible information representation in frontoparietal cortex is not limited to the SR mapping stage, but applies across the functional stages of cognitive processing, thus maximizing neural efficiency and supporting flexible cognition.

---

Goal-directed navigation of complex tasks is key to life outcomes, including longevity (Gottfredson & Deary, 2004) and academic performance (Blair & Razza, 2007). Nevertheless, the neural mechanisms that underlie complex goal-directed cognition are not well understood (Botvinick & Cohen, 2014; Braver, 2012). Whereas distinct regions and networks are critical to domain-specific cognition (e.g. attention; Scolari, Seidl-Rathkopf, & Kastner, 2015; Szczepanski & Knight, 2014), activity in parts of frontoparietal cortex shows overlap across a wide range of tasks and domains (Cole et al., 2013; Duncan, 2010; Hampshire & Sharp, 2015).

Regions of frontoparietal cortex that increase activation across cognitive domains and tasks have been referred to as multiple demand (MD) regions (Duncan, 2010), flexible cognitive hubs (Cole et al., 2013; Cole & Schneider, 2007), and task-activation cortex (Hampshire & Sharp, 2015; Seeley et al., 2007). These mid-dorsolateral, -ventrolateral, and dorsal anterior cingulate regions process information critical to goal-attainment (Fedorenko, Duncan, & Kanwisher, 2013). Specifically, they seem to flexibly represent a different type of information (e.g. stimulus shape, size, or relevance) across different tasks (Crittenden, 2016; Duncan, 2010).

Nevertheless, there is evidence that MD regions may not be homogeneous (Shenhav et al., 2013). Crittenden et al. (2016) manipulated the (complexity of) response rules that had to be learned and found that one subset of MD regions - dorsolateral PFC, inferior frontal sulcus (IFS), and inferior parietal sulcus (IPS) -showed strong representations of task rules. Another subset of MD regions -anterior insula (AI), anterior prefrontal cortex (PFC), and dorsal anterior cingulate cortex (ACC) - showed weaker rule representation. In contrast to the findings by Crittenden et al. (2016), Woolgar et al. (2015) demonstrated that nearly all MD regions represented stimulus-to-response mapping task rules (Woolgar, Afshar, Williams, & Rich, 2015). Thus, these two studies provide conflicting answers as to whether MD regions are functionally homogeneous, even though both studies used multi-voxel pattern analyses (MVPA) and assessed stimulus-response (SR) processes. Therefore, the current study further investigated functional homogeneity across MD regions.

Critically, both studies above investigated the representation of SR-type information in MD regions. However, complex goal-directed cognition requires a series of partially serial, partially parallel functional stages of processing. Indeed such a logical sequence of operations – Target Detection, Stimulus-Response (SR) mapping, Response Execution – forms the core of many long-standing models of cognitive processing (Hoffman, 1975; Pashler, 1984; Stanovich & Pachella, 1977; Yantis & Jonides, 1984; Welford, 1952). The aim of the current study was to assess whether MD regions flexibly represent information across different stages of cognitive processing. So far studies have shown comparable MD activity across cognitive domains, but a strong claim of flexibility also requires MD regions to flexibly represent target selection or SR mapping information depending on current task demands. Moreover, we assessed whether some MD regions more readily represent target selection or SR stage information, but nevertheless represent information related to the other stage if needed.

We developed a search task in which demands on target selection and the SR mapping were manipulated independently under identical visual input. Consequently, differences in brain activity could arise only from differences in complexity of the target selection or SR mapping. Subjects were presented with search arrays and had to select two targets (easy or difficult), apply an SR mapping rule (easy or difficult), and execute the response. Easy target detection consisted of a pop-out search, whereas difficult detection required effortful search. Easy SR mapping consisted of a single-component mapping rule, whereas difficult mapping consisted of two components. We used MVPA because of its sensitivity in discriminating between experimental conditions (Norman, Polyn, Detre, & Haxby, 2006) and employed a well-established functional connectivity analysis to assess MD connectivity changes (Friston et al., 1997; O’Reilly, Woolrich, Behrens, Smith, & Johansen-Berg, 2012). We also employed a whole-brain univariate analysis to assess directional amplitude involvement of non-MD regions in selection and SR mapping.

## Materials & Methods

### Participants

Eighteen healthy, right-handed participants (16 females, mean ± SD age, 24 ± 3.2 years) completed the study. They provided informed written consent and were reimbursed for their time ($20/1.5h). Of the twenty-three participants that took part in the study, data from 2 participants was excluded due to excessive head movement (>10 mm translation and/or 6° rotation) and for another 3 due to behavioral accuracies being below our a-priori criterion (70% correct). The study was approved by the University of Queensland Human Research Ethics Committee.

### Experimental task and procedure

Participants were scanned while performing a visual search task that included a target selection, SR mapping, and response execution operation (Figure 1A). On each trial, subjects were required to, 1) select two targets stimuli, 2) apply the correct SR rule to map the targets onto the correct response, and 3) execute a key press. The task was organized in miniblocks of four trials. At the start of each miniblock, target selection and SR instructions were displayed on the screen for 6s, and followed by a sequence of 4 trials consisting of fixation for 1s, search display for 1.5s, and a blank screen for 1.5s. Responses were recorded from the onset of the search display until the offset of the blank screen (3s). Critically, difficulty of target selection and SR mapping were independently manipulated whilst visual input was identical across conditions.

**Figure 1.**
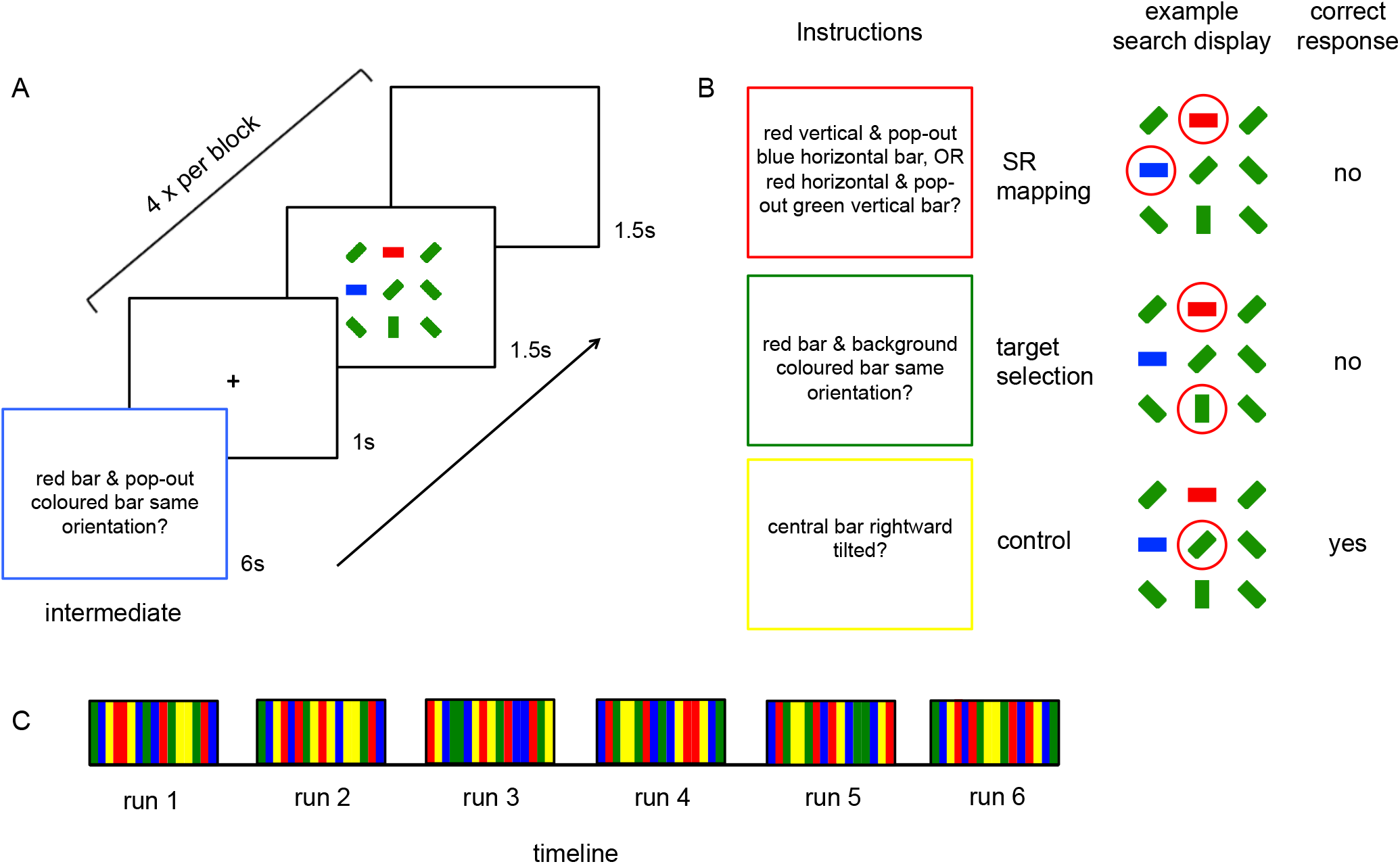
The visual search task was conducted using a block design with four main conditions: control, intermediate, SR, and selection. A, A mini-block consisted of task instructions and four search trials. Displayed are instructions for the intermediate condition (‘do the two pop-out colored targets have the same orientation?’). B, The shortened instructions for the SR (‘are there a red, horizontal pop-out colored and a green vertical pop-out colored bar OR are there a red, vertical pop-out colored and a blue, horizontal pop-out colored bar?’), selection (‘are there a red pop-out colored and a green or blue background-colored bar of the same orientation?’), and control (‘is the central bar tilted rightward?’). The targets in the example search display on the right are encircled (red) for each condition. The correct answer in each condition for the example search display is presented rightmost. Colored instruction borders and red target circles were added to the figure for clarity, but not present in the experiment. C, The experimental time course is depicted as an alternating block design with each run consisting of 16 interleaved blocks (4 of each condition; blue = control, yellow = intermediate, red = SR, green = selection) according to a Latin-square procedure.

### Apparatus & stimuli

The task was created using the Psychophysics Toolbox (version 3.0.8) (Brainard, 1997; Pelli, 1997) for MATLAB (MathWorks, release 2012b). Responses were recorded with an MR-compatible fiber-optic 2-button response box and stimuli were projected onto a screen at the top-end of the scanner bore that was viewed through a mirror mounted to the head coil.

We created 96 search displays that consisted of 9 bars arranged in a 3 × 3 grid and subtended a visual angle of ∼3° (see Figure 1A and 1B). Six bars in each display were tilted at a 45° angle (three left- and rightward each) and all colored green or blue with equal probability. The three bars in each display were oriented horizontally or vertically (equiprobably^1^) and colored red (RGB values = 247,0,0), green (24,145,0), and blue (0,50,254) respectively. Across search displays, color, orientation, and location were controlled. Stimuli were rendered on a light gray background.

### Experimental conditions

Task-difficulty was manipulated independently for target selection and SR mapping, resulting in easy vs. difficult conditions. For the easier SR mapping, the correct response was contingent on a single component SR mapping rule (orientation). For the more difficult SR mapping, the response was contingent on a double component SR mapping rule (orientation and color). For the easier target selection, both target(s) popped-out of the display because of their distinct color (e.g. red and blue amongst green distractors; see Figure 1A and 1B, top row). For the more difficult target selection, one target popped-out (red) but the other was distractor-colored and thus required effortful search (see red and green target in Figure 1B, middle row). Finally, in a control condition, the selection and SR demands were minimal, as participants had to determine whether a central bar was tilted left- or right-ward (in identical displays).

In the fMRI experiment we included four conditions (1) difficult SR / easy selection (‘are there a red, horizontal pop-out colored and a green vertical pop-out colored bar OR are there a red, vertical pop-out colored and a blue, horizontal pop-out colored bar?’), henceforth referred to as *SR (mapping)*, (2) difficult target selection / easy SR (‘are there a red pop-out colored and a green or blue background-colored bar of the same orientation?’), henceforth referred to as (target) *selection* (3) easy SR and easy selection (‘are the two pop-out colored bars of the same orientation?), henceforth referred to as *intermediate*, and (4) a control condition with minimal SR and selection requirements (‘is the central bar tilted rightward?’), henceforth referred to as control.

### Procedure

The four conditions were interleaved using mini blocks (22s). Sixteen blocks (four of each condition) were interleaved according to a Latin-square procedure in each run for six runs (Figure 1 panel C). Each visual search display was shown once per condition and display order was determined by a random permutation of all displays. The total number of trials was 384 and the scanning session lasted about 50 minutes. Before transferring into the scanner participants completed a training block of 96 trials (24 of each condition; 15minutes).^2^

### fMRI acquisition

Anatomical and functional images were acquired using a 3T Siemens Trio MRI scanner (Erlangen, Germany) and a 32-channel head coil. Functional T2*-weighted images were acquired parallel to the AC–PC plane using a GRE (gradient recalled echo) EPI (echo planar imaging) sequence (TR = 1.8s, TE = 30ms, FA = 80°, FOV = 192 × 192, matrix = 64 × 64, in-plane resolution = 3 × 3 mm). Each volume consisted of 29 slices with a thickness of 3 mm and a 0.3-mm inter-slice gap. At the start of the session, we collected a T1-weighted anatomical image using an MP-RAGE sequence (TR = 1.9s, TE = 2.32ms, FA = 9°, FOV = 230 × 230 × 230, resolution = 0.9mm^3^). Each run consisted of 196 volumes. The first 3 images of each run were discarded to account for T1 equilibration effects.

### fMRI preprocessing

Images were preprocessed and analyzed with BrainVoyager (version 20.2; Brain Innovation, Maastricht, The Netherlands) and custom-written MATLAB code using the BVQX Toolbox v08d (Weber, 2014). Images were slice-scan time corrected, re-aligned to the first image, and coregistered with the structural images. Finally, data were normalized to the standard Montreal Neurological Institute (MNI) template. For the univariate analyses and connectivity (PPI) analyses, data were smoothed with an 8mm full-width-half-maximum Gaussian kernel. For the MVPA, unsmoothed data were used.

### ROI selection

To minimize any theory-driven bias in ROI selection, a single run from each participant’s data was selected (equiprobably from runs 1-6) to compute a functional localizer contrast (this run was excluded from subsequent MVPA and connectivity analyses). The localizer contrast showed voxel clusters (minimum size of 10 voxels) where BOLD activity averaged over the SR, selection, and intermediate conditions exceeded that in the control condition (FDR q: .01). The contrast yielded twelve activation clusters, all in frontal and parietal cortex. We computed the centre of gravity coordinates for each cluster (see Table 1, grey highlighted regions/coordinates) and created a spherical ROI with a radius of 6mm around each.^3^

The majority of identified ROIs were similar to those identified in previous studies (Dosenbach et al., 2007; Duncan, 2010)(See table 1, ROIs marked ++). We included an additional 4 ROIs based on MD coordinates reported previously (Duncan, 2010) that were not covered by the twelve ROIs we identified with the functional localizer. Together these sixteen ROIs covered the MD regions that were studied in previous research and our procedure ensured that other regions with significant task-activation were not excluded on a-priori theoretical grounds.

**Table 1.** ROIs.

| ROI name | ROI coordinates |  |  | t-value<br>(FDR-corrected) | Duncan<br>(2010) |  |  | Dosenbach<br>(2007) |  |  |
| --- | --- | --- | --- | --- | --- | --- | --- | --- | --- | --- |
|  | x | y | z |  | x | y | z | x | y | z |
| **L AI / FO <sup>++</sup>** | -32 | 21 | 3 | 6.158 | -35 | 18 | 2 | -35 | 14 | 5 |
| **R AI / FO <sup>++</sup>** | 32 | 23 | 0 | 7.158 | 35 | 18 | 2 | 36 | 16 | 4 |
| **ACC <sup>+</sup>** | 0 | 31 | 24 | — | 0 | 31 | 22 | — | — | — |
| **L MFG** | -24 | 4 | 52 | 7.303 | — | — | — | — | — | — |
| **R MFG** | 24 | 4 | 52 | 5.843 | — | — | — | — | — | — |
| **L IPS <sup>++</sup>** | -31 | -50 | 48 | 10.686 | -37 | -56 | 41 | -31 | -59 | 42 |
| **R IPS <sup>++</sup>** | 31 | -49 | 48 | 9.176 | 37 | 56 | 41 | 30 | -61 | 39 |
| **L rPFC <sup>+</sup>** | -21 | 43 | -10 | — | -21 | 43 | -10 | -25 | -44 | -12 |
| **R rPFC <sup>+</sup>** | 21 | 43 | -10 | — | 21 | 43 | -10 | 25 | -44 | -12 |
| **L IFS (dlPFC) <sup>++</sup>** | -46 | 32 | 29 | 4.474 | -41 | 23 | 29 | -43 | 22 | 34 |
| **R IFS (dlPFC) <sup>++</sup>** | 42 | 39 | 22 | 4.595 | 41 | 23 | 29 | 43 | 22 | 34 |
| **L FC <sup>++</sup>** | -43 | 5 | 30 | 3.891 | — | — | — | -41 | 3 | 36 |
| **R FC <sup>++</sup>** | 48 | 9 | 28 | 6.720 | — | — | — | 41 | 3 | 36 |
| **Pre SMA <sup>+</sup>** | 0 | 18 | 50 | — | 0 | 18 | 50 | — | — | — |
| **L precuneus** | -29 | -71 | 34 | 10.686 | — | — | — | -9 | -72 | 37 |
| **R precuneus** | 30 | -65 | 35 | 9.176 | — | — | — | 10 | 69 | 39 |

<sup>++</sup> ROIs identified in our functional localizer that were similar to ROIs identified in Duncan, 2010 or Dosenbach, 2007. <sup>+</sup> ROIs that were identified in Duncan, 2010, but that failed to reach significance in our functional localizer.

### Analyses

In line with previous studies (Woolgar et al., 2015) we used MVPA to assess condition differences in the pattern of activity across voxels because of its sensitivity in discriminating between experimental conditions (Norman et al., 2006). Subsequently, we employed a well-established functional connectivity analysis (Friston et al., 1997; O’Reilly et al., 2012) to assess whether multi-voxel activity pattern differences across the selection and SR manipulation were accompanied by a differential pattern of connectivity between MD regions. Finally, we conducted a whole-brain univariate analysis to assess the directional amplitude of the effects and involvement of non-MD regions in selection and SR mapping.

#### MVPA analyses

MVPA was performed using BrainVoyager (version 20.2; Brain Innovation, Maastricht, The Netherlands) to assess whether voxel activity patterns for different levels of complexity in the target selection and SR mapping stages were distinguishable. In a first step, GLM analyses were performed on each individual’s data (5 runs), with separate regressors for each condition. Each regressor was modeled as a rectangular function from the onset of the first search display in each block to the end of the fourth trial and convolved with the canonical HRF. We derived t-values for each voxel in each ROI per mini-block. Values were z-scored across all voxels to reduce any impact of small differences in univariate activity between selection and SR conditions on MVPA results.

The pattern discrimination between tasks was then estimated using pairwise classification between SR and control, and selection and control for each participant. We used a support vector machine (SVM) with a linear kernel and optimized cost parameter C in a level-split procedure. The classifier was trained on 4 runs and tested on one. This procedure was repeated five times on a leave-one-out basis (fivefold cross-validation) such that each run was used for testing once. Classification accuracies for each ROI were averaged across splits and participants to yield a single classification accuracy.

To assess whether classification accuracy was significantly higher than chance, we ran 200 permutation tests per split with condition labels randomly shuffled, and identified the 95^th^ percentile of this distribution. Then we assessed whether pattern discrimination with intact labels exceeded the observed 95^th^ percentile of this distribution. Finally, classification accuracy for SR and selection were compared by a series of paired two-sided t-tests.

It has been noted that RT differences may confound task-related influences on neural activity (for an MVPA-specific discussion see e.g. Todd, Nystrom, & Cohen, 2013; Woolgar, Golland, & Bode, 2014). Therefore, we assessed the influence of cross-condition RT differences (SR vs. control and selection vs. control) on classification accuracy for the respective comparisons (identical approach to Crittenden et al., 2016; Erez & Duncan, 2015). For each condition pair, we conducted a regression analysis of classification accuracy on the RT difference between the two respective conditions (SR vs. control and selection vs. control). In a first step, we extracted, for each participant, the classification accuracy for each condition pair in each ROI. In a second step, we computed an average classification accuracy across ROIs for each participant. In a third step, for each participant we computed the RT difference for each condition pair (i.e. RT SR minus control; RT selection minus control). We then regressed classification accuracy on RT for each subject and extracted the slopes (β) and intercept (α). The slope parameter should be positive and significantly greater than 0 if RT significantly modulates classification accuracy and if classification could be explained entirely by RT then one could expect that at the intercept (no RT difference) the classification accuracy should be at chance-level (50%).

#### Connectivity analyses

Connectivity analyses were performed using a standard psychophysiological interaction analysis (PPI; Friston et al., 1997) for the contrast SR versus selection with the PPI plugin for BrainVoyager. The preprocessed and smoothed data were used for this analysis. The source regions assessed were those ROIs that showed a difference in classification accuracy in the MVPA analysis for SR and selection. The potential receiving voxels were restricted to the ROIs introduced in Table 1. Connectivity strength was calculated across mini blocks.

First we extracted the time series from the first-level GLM for each source ROI and convolved it with the vector that represents the contrast SR versus selection. Subsequently we ran a second-level GLM with all three task difficulty variables and looked at the unique explanatory power of the PPI. Changes in connectivity between SR and selection are evident when PPI regressor values differ significantly from zero (FDR q: .01, minimum cluster size 20 voxels).

#### Whole-brain univariate analyses

Finally, to test for activations uniquely associated with either target selection or SR mapping in other brain areas, we also conducted a standard whole-brain analysis with the contrasts SR – intermediate and selection - intermediate (threshold of p-FDR < 0.01; 15 continuous voxels).

## Results

### Behavioral results

Mean reaction times (RTs) differed across conditions, *F*(3,17) = 152.02, *p* < .001, *partial η^2^* = .96 (see Figure 2). Paired, two-tailed t-tests showed that RTs in all conditions differed from one another with RTs in the control condition being faster than in the intermediate condition (*t*(19) = 10.89, *p* < .001), which in turn were faster than those in the SR condition (*t*(19) = 9.78, *p* < .001), which were faster than those in the selection condition (*t*(19) = 3.10, *p* < .001). Mean error percentages are presented in Figure 2 as well (black circles). There was no evidence of a speed-accuracy trade-off.

**Figure 2.**
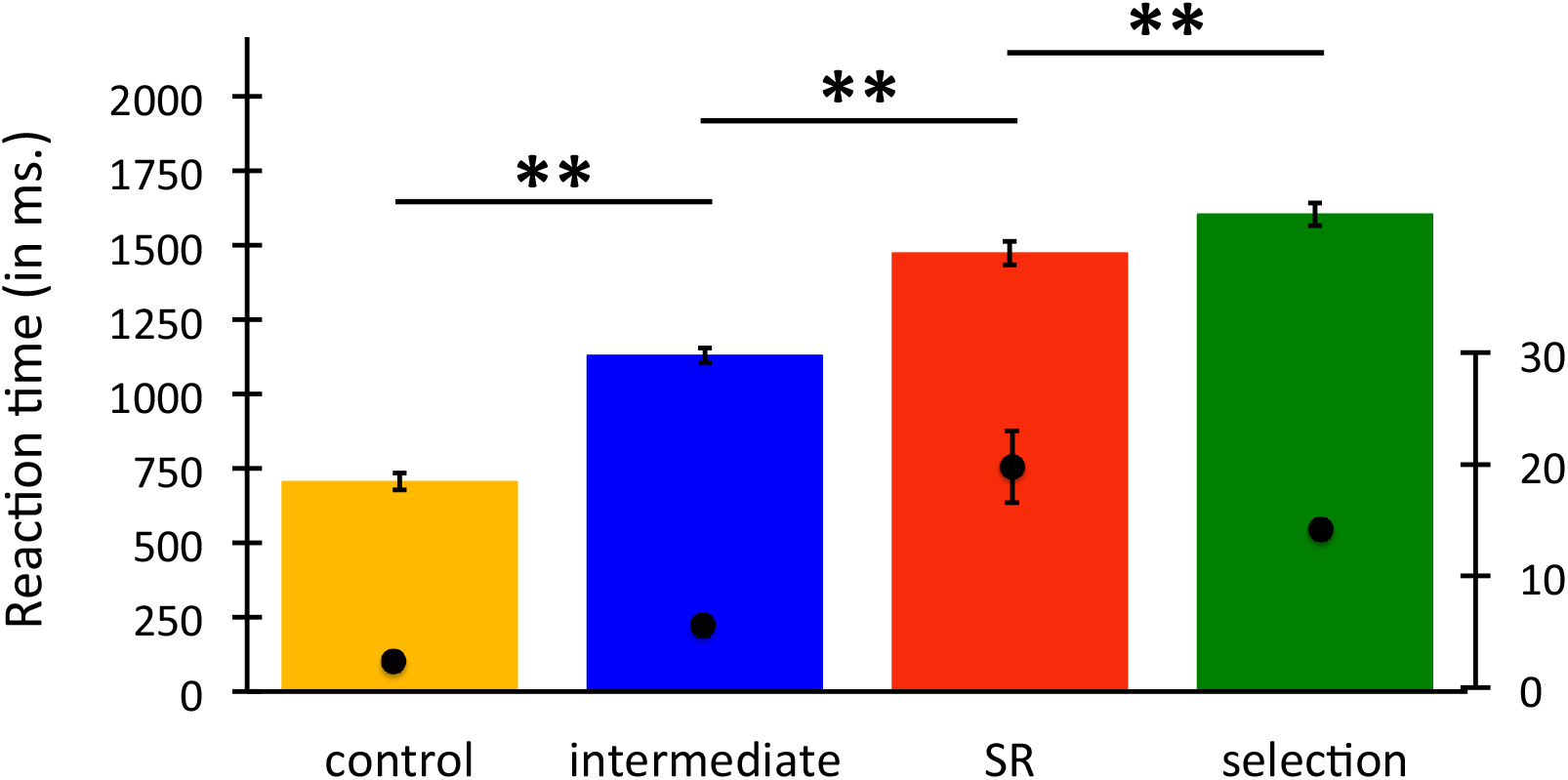
Mean RTs (bars) differed significantly between all conditions (control < intermediate < SR < selection). Mean error percentages are plotted as black circles. Error bars represent the standard errors of the means and may be smaller than the plotting symbol. ** *p* < 0.001.

### MVPA results

The majority of ROIs showed significant decoding of task difficulty (for SR vs. control as well as selection vs. control) with the average classification accuracy of increased task difficulty over all ROIs being 75%. Figure 3 depicts the mean classification accuracy for SR vs. control (red bars) and selection vs. control (green bars) and the 95^th^ percentile of the null distribution (grey line) that was used as the criterion for significance. Whereas nearly all regions decoded both difficulty of the target selection as well as difficulty of the SR mapping, some regions showed significantly stronger decoding for one or the other.

To compare classification accuracy for target selection and SR, we conducted a repeated- measures ANOVA with cognitive stage of processing (selection vs. SR) and ROI as within-groups factors. We observed a significant effect of ROI (*F*(8,9) = 4.642, *p* = 0.017) and a significant interaction of cognitive stage of processing and ROI (*F*(8,9) = 4.199, *p* = 0.016). A series of follow-up t-tests showed that, a first set of ROIs (AI/FO and ACC) showed comparatively stronger decoding for the task pair that differed in the SR dimension (see Figure 3 left cluster). For the AI/FO classification accuracy was 77% (SR) vs. 70% (selection), *p* = .009. For the ACC classification accuracy was 74% (SR) vs, 67% (selection), *p* = 0.013. A second set of ROIs (MFG and IPS) showed the opposite pattern with comparatively stronger decoding for the task pair that differed in the selection dimension (Figure 3, central cluster). For the MFG classification accuracy was 71% (SR) vs. 77% (selection), *p* = .010. For the IPS classification accuracy was 74% vs 81%, *p* = 0.007. For the precuneus classification accuracy was 76% vs 81%, *p* = 0.038. The remaining ROIs (IFS, FC, pre-SMA, and precuneus) did not show a significant difference in decoding strength for SR and selection (see Figure 3, right cluster; IFS: 77% vs. 76%, *p* = .595; FC: 76 vs. 79%, *p* = 0.184; pre-SMA: 76 vs. 76%, *p* = .667; rPFC: *p* = .470). The only frontoparietal ROI that did not show significant decoding for either task pair was the rPFC.

**Figure 3.**
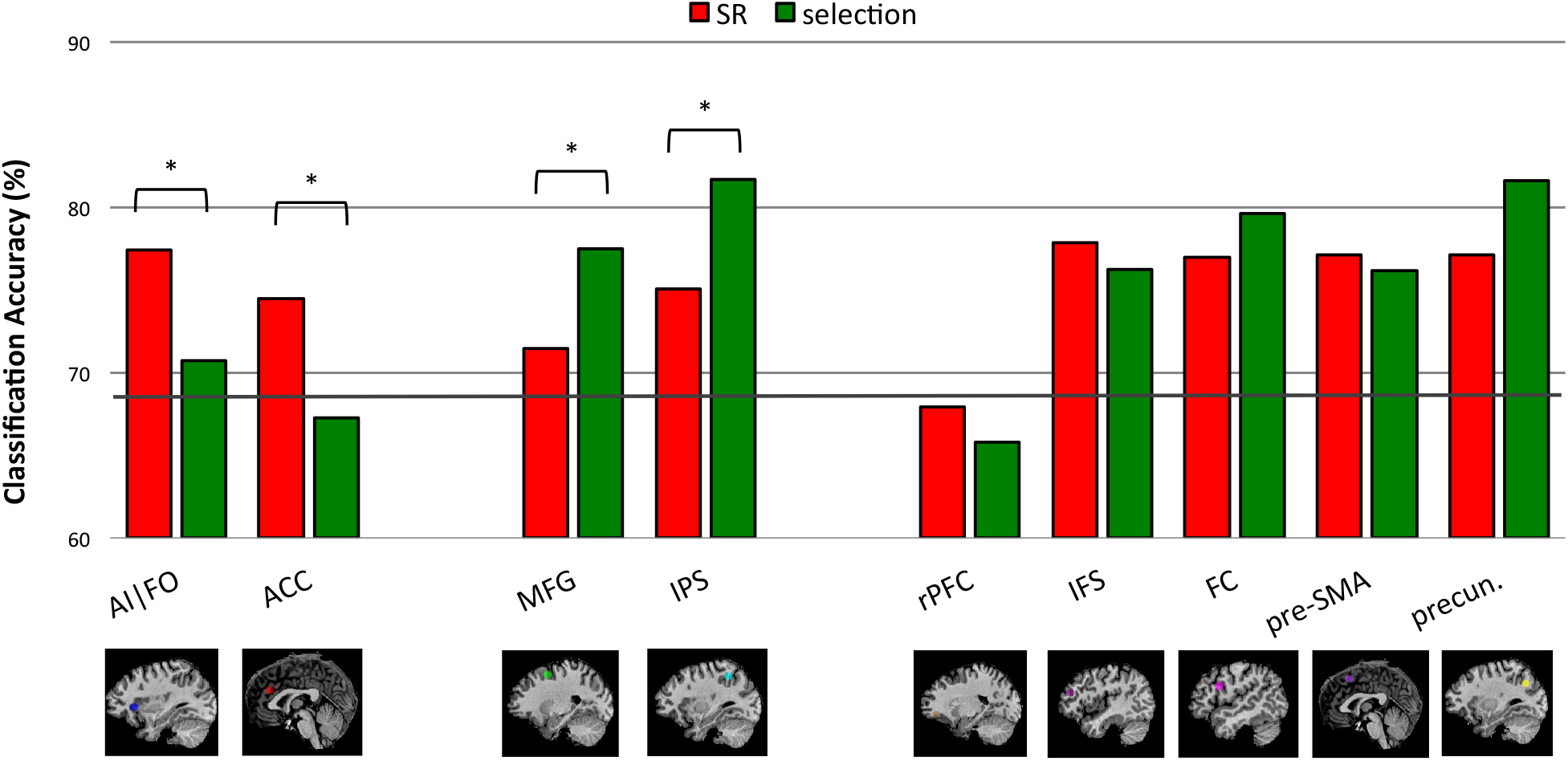
Task decoding in selected frontal and parietal ROIs. Red bars represent task decoding for the difficult SR vs control (easy SR) conditions; green bars represent task decoding for the difficult selection vs control (easy selection) conditions. ROIs with comparably stronger decoding of SR-related information are shown on the left, ROIs with significantly stronger decoding of target-selection-related information are shown in the middle, and ROIs that show comparable decoding strength for SR and selection are displayed on the right. The grey line denotes the 95^th^ percentile of the chance distribution. 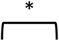 denotes a significant difference (corrected p value < .05) in the pairwise comparison between SR and selection classification accuracy.

To assess whether RT differences between conditions may drive our MVPA results, we then regressed classification accuracy on RT for each subject and extracted the slopes (β) and intercept (α). Critically, classification accuracy at intercept – i.e. in the absence of any RT differences between conditions - was above chance level for SR and selection (Figure 4A). Specifically, for the SR-control and selection-control pairs, the classification accuracy at intercept was 64% and 63% respectively, which two-tailed, Wilcoxon signed rank tests showed to be significantly greater than chance level (ps < 0.05). This indicates task decoding in the absence of RT differences.

The regression slopes provided evidence that RT moderately, but significantly modulated classification accuracy for target selection (Figure 4B). Specifically, For the SR-control and selection-control pairs, the regression slopes were 0.014 and 0.013 respectively. The former did not differ significantly from 0 (*p* = 0.103), whereas the latter did (*p* = 0.024). Since the mean RT difference between our two conditions of interest was around 100ms, this suggests that RT differences may have inflated the classification accuracy difference by ∼1.3%. Therefore, although it seems that RT provides a small contribution to classification accuracy in the current study, such an influence is far from accounting for the full observed effects.

**Figure 4.**
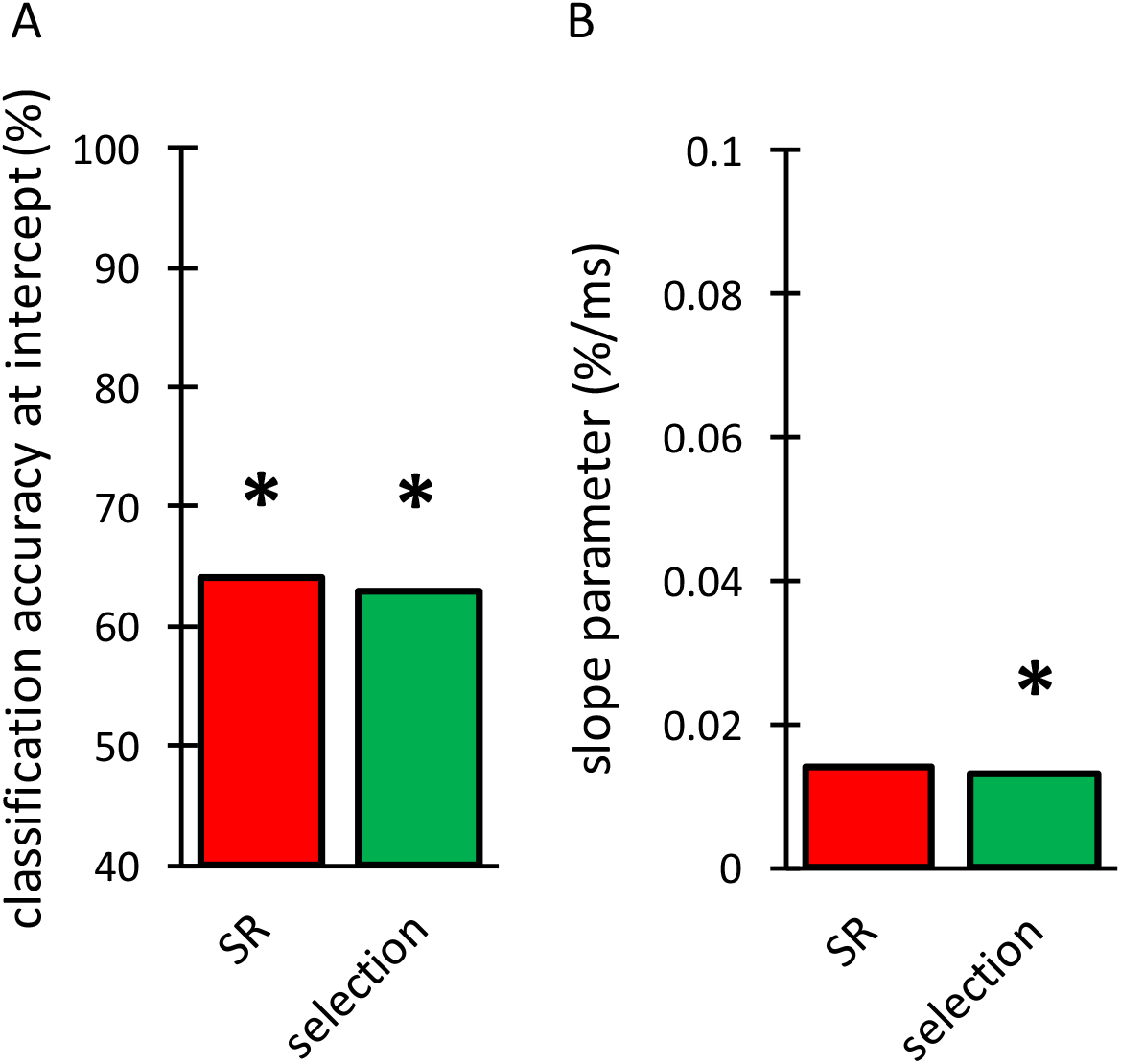
Mean parameter estimates of α and β after regression of classification accuracy differences (SR minus control and search minus control) on absolute response time differences. A, At the intercept (i.e. in the absence of RT differences) the mean value of α for SR (red) and selection (green) was significantly greater than chance (50%). B, Mean value of β for SR and search with only the latter being significantly greater than 0. * p < 0.05.

### PPI results

The PPI analysis showed a change in functional connectivity between several ROIs for the SR and selection condition (see figure 5). Specifically, connectivity between the ACC and right MFG (1167 voxels significant, *t*(17)= 2.783, corrected *p* = 0.015) was stronger during SR and weaker during selection. Similarly, connectivity between the ACC and FC (left FC: 1309 voxels significant, *t*(17)= 3.075, corrected *p* = 0.008; right FC: 1309 voxels significant, *t*(17)= 2.632, corrected *p* = 0.019) was stronger during SR and reduced during selection. The opposite pattern with relatively stronger functional connectivity during selection and reduced connectivity during SR was observed for the FC and the right precuneus (1238 voxels, *t*(17)= -2.786, corrected *p* = 0.016).

**Figure 5.**
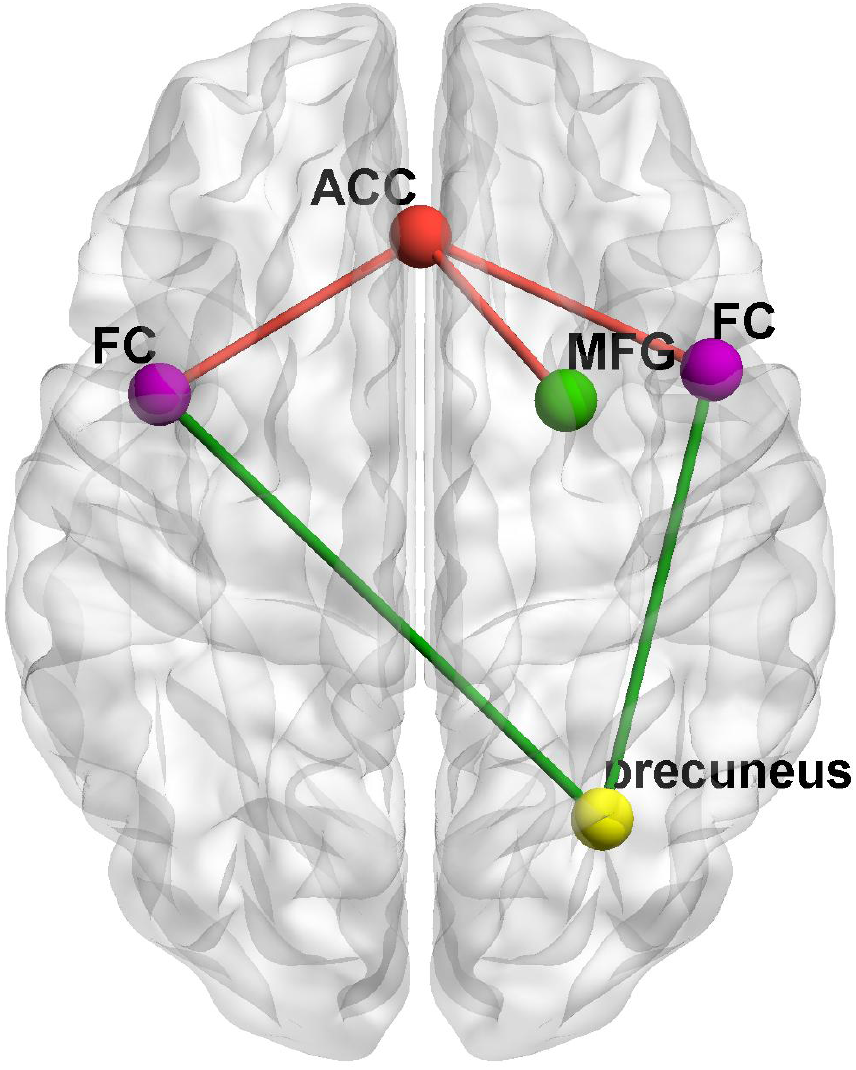
Increased functional connectivity between ROIs during target selection (green lines) and SR (red lines). For target selection, an increased connectivity was observed bilaterally between FC and the right precuneus. For SR, increased connectivity was observed bilaterally between ACC and FC, as well as ACC and right MFG. ACC = red, MFG= green, FC = purple, precuneus = yellow. ROIs and connections colored red are preferentially involved in SR mapping; ROIs and connections colored green are preferentially involved in selection.

### Whole-brain univariate analyses

For target selection (difficult target selection > easy), the whole-brain univariate analysis revealed 4 regions of gray matter activation (see Fig. 6), centered bilaterally on the precuneus (left: -21, -55, 52; 522 voxels; right: 18, -55, 52; 1906 voxels; Brodmann area 7) and the middle occipital gyri (left: -33, -88, 4; 3788 voxels; right: 30, -91, 10; 7179 voxels; Brodmann areas 19).

**Figure 6.**
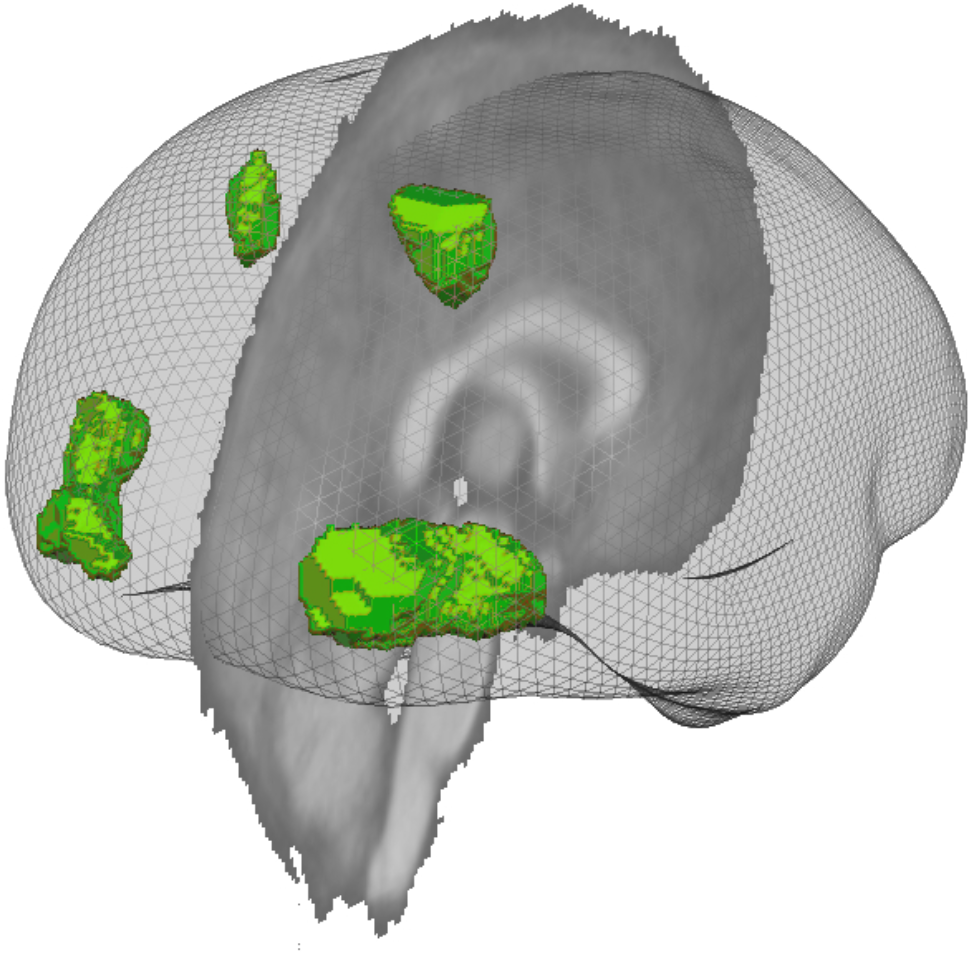
For target selection (difficult target selection > easy), the whole-brain univariate analysis revealed 4 regions of gray matter activation, centered bilaterally on the precuneus (left: -21, -55, 52; right: 18, -55, 52) and the middle occipital gyri (left: -33, -88, 4; right: 30, -91, 10). Figure 5 shows the brain from a posterior viewpoint.

For SR (difficult SR > easy), the whole-brain univariate analysis revealed 8 regions of gray matter activation (see Fig. 7), in frontal, parietal and occipital cortex. Frontal activation included the bilateral anterior middle frontal gyri (left: -37, 59, -11; 12850 voxels; right: 48, 50, -20; 12665 voxels; Brodmann areas 10/11), a more posterior portion of the left middle frontal gyrus (-36, 23, 31; 2340 voxels; Brodmann area 9), the medial frontal gyrus (6, 35, 43; 19952 voxels; Brodmann area 8), and the right claustrum (30, 23, -5; 1376 voxels). Parietal and occipital activation included the bilateral inferior parietal lobule (left: -37, 64, 46; 2021 voxels; right: 48, -55, 43; 9986 voxels; Brodmann areas 7/40), and the cuneus (-6, -70, 31; 6187 voxels; Brodmann area 7). Overall, then, the whole-brain analysis confirmed the pattern of results observed in previous studies, implicating parietal and occipital cortex in target selection and frontal and parietal regions SR.

**Figure 7.**
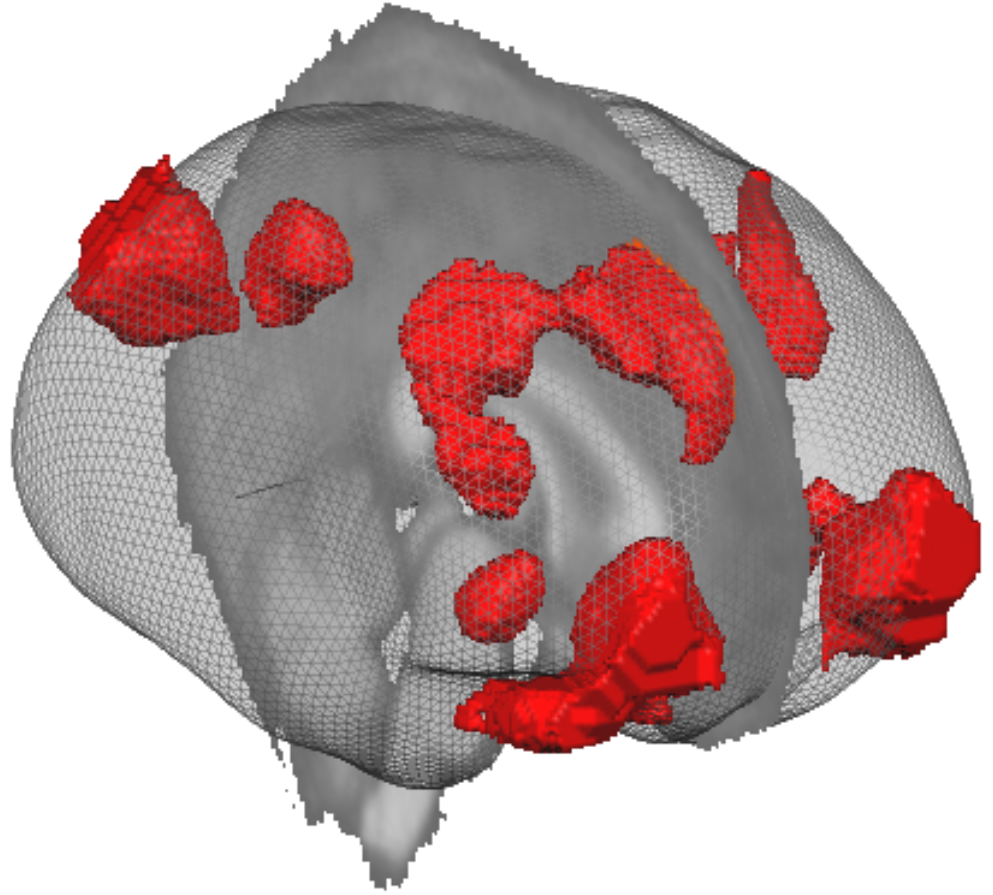
For SR (difficult SR > easy), the whole-brain univariate analysis revealed 8 regions of gray matter activation in frontal, parietal and occipital cortex. Frontal activation included the bilateral anterior middle frontal gyri (left: -37, 59, -11; right: 48, 50, -20), the left posterior middle frontal gyrus (-36, 23, 31), the medial frontal gyrus (6, 35, 43), and the right claustrum (30, 23, -5). Parietal and occipital activation included the bilateral inferior parietal lobule (left: -37, 64, 46; right: 48, -55, 43), and the cuneus (-6, -70, 31). Figure 7 shows the brain from an anterior viewpoint.

## Discussion

Our study is the first to demonstrate that MD regions flexibly represent information from different stages of cognitive processing depending on the current task demands. Whereas evidence of comparable activity in MD regions across a wide range of cognitive domains (e.g. attention, working memory) is critical (Cole et al., 2013; Cole & Schneider, 2007; Crittenden et al., 2016; Duncan, 2010; Hampshire & Sharp, 2015; Seeley et al., 2007), a strong claim of flexible processing in MD regions requires comparable activity across different stages of cognitive processing (McClelland, 1979; Pashler, 1984; Stanovich & Pachella, 1977; Sternberg, 1969; Townsend, 1974; Turvey, 1973). Flexibility of information representation was supported by the observation that nearly all MD regions represented target selection related information when target selection was demanding and SR mapping information when the SR stage of processing was demanding.

Furthermore, the current study shows that some MD regions show relatively stronger representations for a specific stage of processing. Specifically, a demanding target selection stage was associated with selectively enhanced information representation in a relatively more dorsal cluster of regions consisting of MFG and IPS, whereas a demanding SR mapping stage was associated with enhanced representation of information in a relatively more ventral cluster consisting of AI/FO and ACC. Importantly, in our study the visual input (i.e. stimulus displays) was identical across conditions and as such differences cannot be attributed to differences in visual input or response output. Crittenden et al. (2016) showed a unidirectional increase in the representation of task rules in a subset of MD regions (dlPFC, IFS, IPS). Here, we demonstrate a cross-over in which specific regions showed enhanced representation of target selection information (MFG and IPS) and other regions of SR information (AI and ACC).

Three explanations can account for the involvement of the same cortical regions, to the same or a different degree, in processing target selection and SR-related information. Theories of adaptive cognitive control have purported that current goals drive which particular stimulus features are coded by the same neuronal populations in frontoparietal cortex at any given time (Fedorenko et al., 2013). Such an adaptive coding at the neuronal level suggests that current task demands drive the tuning profile of neurons in these regions through changes in the baseline excitability and altered short-term synaptic plasticity (Stokes et al., 2013) or frequency-specific coherence (Buschman, Denovellis, Diogo, Bullock, & Miller, 2012). An as such one may be tempted to interpret our findings as suggestive of neurons flexibly updating whether they code target selection- or SR-related information, whilst simultaneously neurons in specific regions are better equipped to process information that is critical to either target selection (e.g. IPS) or SR mapping (e.g. ACC).

Primate research (Kadohisa et al., 2013; Stokes et al., 2013) has provided some evidence for adaptive coding. However, a single MRI voxel contains more than an estimated one million neurons (Logothetis, 2008). Therefore, fMRI studies cannot disentangle adaptive coding from the alternative explanation that MD regions have a neuronal organization where adjacent neurons respond to different visual properties. This would be much akin to the organization in early visual cortex where one neuron is sensitive to edge orientation whilst a neighboring neuron can be sensitive to e.g. motion direction or spatial frequency. Neurons that are critical to successful target detection may be closely interspersed with neurons that are critical to SR mapping. The current task goal and task demands may determine which particular subpopulation of neurons in a region becomes responsive and different subpopulations may well fall within the same MRI voxel and thus occur at a scale too small to be detected by MRI.

Another explanation for comparable activity across stages of processing is that these MD regions may be involved in a singular higher-level cognitive process that operates during different stages of cognitive processing. For example, previous research has suggested that frontoparietal cortex may be involved in the creation of subtasks for goal-achievement (Farooqui, Mitchell, Thompson, & Duncan, 2012) or the monitoring of conflict and the resulting adjustment in the level of effortful cognitive control (Botvinick & Braver, 2015; Botvinick & Cohen, 2014; Botvinick, Cohen, & Carter, 2004; Shenhav, Botvinick, & Cohen, 2013). A conflict monitoring role could explain why frontoparietal regions (especially the ACC) become more active in a wide range of SR tasks as difficulty increases (Crittenden et al., 2016; Woolgar et al., 2015). Differences in coding accuracy in the current study may be argued to reflect differences in the need for conflict monitoring: e.g. the SR mapping manipulation may be argued to require more cognitive control because it was more complex than the difficult target selection (as indicated by a slightly elevated error rate). Alternatively, the ACC may particularly be involved in processing conflict at the SR stage and not (or less) at the target selection stage. Increased involvement of the ACC in SR mapping is in line with previous research implying this region represents and maintains current goals at an abstract, task-independent level (Lopez-Garcia et al., 2015; Munakata et al., 2011), and the proposal that the dlPFC/ACC monitors goal progress (Benn et al., 2014). However, such a singular process that would be active across both the target selection and SR mapping manipulations cannot account for the current results as we observed changes in connectivity as well as changes in region-specific information representation in e.g. the ACC and precuneus.

Particularly, the ACC showed selectively enhanced coding of SR related information and this was accompanied by an increase in connectivity between the ACC and several other regions (right MFG and FC) during the SR stage. Conversely during the target selection stage of task processing, selectively enhanced coding of information by MFG was accompanied by a decrease in MFG-ACC connectivity as well as a decrease in connectivity between FC and ACC, but an increased connectivity between FC and right precuneus. Overall, such changes in connectivity render it less likely that changes in coding performance simply reflect on an identical operation that is activated to different degrees as this should draw on a comparable patterns of functional connectivity.

Our study extends previous research that has suggested that frontoparietal cortex includes several regions that flexibly change their functional connectivity based on the demands of the task at hand (Cole et al., 2013). Flexible hub theory proposes two specific systems-level neural mechanisms that support adaptive task control. Global variable connectivity refers to the fact that several regions in frontoparietal cortex show different functional connectivity patterns across tasks. Compositional coding suggests that such shifts in connectivity may be systematic and thus may be reused and recombined across different domains and tasks to enable transfer across tasks. Together, these two mechanisms describe a distributed coding system that provides an efficient means of implementing a wide variety of task states. We propose that compositional coding may be guided by the stages of cognitive processing, such that there is a systematic relationship between connectivity patterns and the stages of cognitive processing. This framework allows well-established connections to be dynamically adjusted according to the current task demands. Perhaps even more importantly, this account allows for the transfer of knowledge and skills to novel tasks.

We, as others have done before us, suggest that our neural system is faced with the need to exert goal-directed control in an efficient yet flexible way such that it is effective across a wide range of tasks and situations. One way to implement such flexible yet efficient cognitive control is to identify the building blocks that are inherent to nearly all tasks. Other studies have pointed to stimulus feature dimensions as such potential building blocks – e.g. size, shape, texture (Botvinick & Cohen, 2014). Our proposal suggests that goal-driven neural activity relies on a sequence of cognitive processing stages: target selection, SR mapping, and response execution.

In sum, frontoparietal resources are flexibly assigned to the functional stages of cognitive processing according to current task demands. At the same time, some flexible frontoparietal regions are relatively more adept at representing information from a specific functional stage. A neurocognitive organization in which some regions of frontoparietal cortex are preferentially sensitive to a specific functional stage of processing and at the same time allocation of resources across functional stages of processing is flexibly driven by stage demands, ensures efficient functioning in an ever-changing environment. Overall, this study provides one approach to look beyond domain specific models to explain flexible neural activity (Cole et al., 2013; Duncan, 2010; Hampshire & Sharp, 2015).

## Footnotes

1 with the restriction that they could not all three be of the same orientation.

2 At the end of each trial, accuracy feedback was provided. If a participant’s accuracy rate was below 70% in any of the four conditions, participants had to complete the training block a second time. The only other divergence in procedure from the scanner block was that instructions were displayed for 12 seconds.

3 Peak voxel coordinates for the single run localizer were representative of all six runs with the distance between the peak coordinates observed for a single run and those observed when using all six runs being less than 3 voxels (x mean: 1.8, y mean: 2.6, z mean: 1.8).

